# *Strongyloides stercoralis*: spatial distribution of a highly prevalent and ubiquitous soil-transmitted helminth in Cambodia

**DOI:** 10.1101/453274

**Authors:** Armelle Forrer, Virak Khieu, Penelope Vounatsou, Paiboon Sithithaworn, Sirowan Ruantip, Rekol Huy, Sinuon Muth, Peter Odermatt

## Abstract

**Background:** *Strongyloides stercoralis* is a neglected soil-transmitted helminth that occurs worldwide and can cause long-lasting and potentially fatal infections due to its ability to replicate within its host. *S. stercoralis* causes gastrointestinal and dermatological morbidity. The objective of this study was to assess the *S. stercoralis* infection risk, and using geostatistical models, to predict its geographical distribution in Cambodia.

**Methodology / Principal Findings:** A nation-wide community-based parasitological survey was conducted among the population aged 6 years and above. *S. stercoralis* was diagnosed using a serological diagnostic test detecting antigens in urine. Data on demography, hygiene and knowledge about helminth infection were collected. *S. stercoralis* prevalence among 7,246 participants with complete data record was 30.5% and ranged across provinces between 10.9% and 48.2%. The parasite was ubiquitous in Cambodia, with prevalence rates below 20% only in five south-eastern provinces. Infection risk increased with age both in men and women although girls aged less than 13 years and women aged 50 years and above had lower odds of infection than their male counterparts. Open defecation was associated with higher odds of infection while declaring having some knowledge about health problems caused by worms was protective. Infection risk was positively associated with night maximum temperature, minimum rainfall, and distance to water, and negatively associated with land occupied by rice fields.

**Conclusions / Significance:** *S. stercoralis* infection is ubiquitous and rampant in Cambodia. The parasite needs to be addressed by control programs delivering ivermectin. However the high cost of this drug in Cambodia currently precludes control implementation. Donations, subsidization or the production of affordable generic production are needed so *S. stercoralis*, which infests almost a third of the Cambodian population, can be addressed by an adequate control program.

**Authors Summary:** The threadworm, *Strongyloides stercoralis*, is a most neglected worm infection transmitted through infective larvae on the soil. Threadworms occur worldwide and particularly in tropical climates. It may cause long-lasting and potentially fatal infections due to its ability to replicate within its host. This study aimed to assess the risk of threadworm infection in at the national level in Cambodia.

We conducted a nation-wide community-based parasitological survey among the population aged 6 years and above. The threadworm was diagnosed using a serological diagnostic test detecting antigens in urine. Data on demography, hygiene and knowledge about helminth infection were collected. The threadworm infection risk was calculated by using geostatistical models to predict its geographical distribution in Cambodia. About one third (30.5%) of the enrolled study participants (n=7,246) were infected with threadworms. The lowest and hightest infection rates a province level was 10.9% and 48.2%, respectively. Prevalence rates below 20% were found only in five south-eastern provinces. The risk of an infection with threadworms increased with age in men and women. Open defecation was associated with higher risk of infection while declaring having some knowledge about health problems caused by worms was protective. Furthermore, the threadworm infection risk was positively associated with environmental factors such as night maximum temperature, minimum rainfall, and distance to water, and negatively associated with land occupied by rice fields.

Threadworm infection is highly prevalent in Cambodia and adequate control measures are warranted, including access to treatment, in order to address the burden of this NTD in Cambodia.

## Introduction

*Strongyloides stercoralis* is a highly neglected intestinal nematode which larvae living in fecally polluted soil infect humans transcutaneously, like hookworms. *S. stercoralis* occurs worldwide but thrives in warm regions with poor sanitation conditions [1]. *S. stercoralis* has been under-detected and overlooked for decades because its larvae are not detected by standard field diagnostic techniques [1–5]. Up to recently, the only available estimates originated from a review conducted in the late 80s estimating that there would be 30-100 million cases worldwide [6]. More recent estimates include prevalence rates between 10% and 40% in subtropical and tropical countries, while, based on the ratio of hookworm *vs. S. stercoralis* cases in studies using diagnosis approaches adequate for the latter, *S. stercoralis* prevalence could be half of hookworm’s, i.e. 200-370 cases worldwide [1, 7, 8].

In Cambodia, *S. stercoralis* was found to be highly prevalent in two community-based large-scale surveys documenting prevalence rates of 25% and 45% in the southern province of Takeo and the northern province of Preah Vihear, respectively [9, 10]. *S. stercoralis* infection is more prevalent among adults due to its unique ability among STH to replicate within the host, which leads to infections that can last for decades in absence of treatment [11]. Importantly, in case of immunosuppression, this auto-infection cycle accelerates and results in hyperinfection, a condition that is 100% fatal if untreated [12–14]. Additionally, chronic infection with *S. stercoralis* may cause abdominal pain, nausea, vomiting, diarrhea, as well as urticaria and larva currens [15–17]. The latter is a serpiginous intermittent moving eruption due to the parasite migration under the skin. Its location on the buttocks, thighs and trunk, together with the high speed of migration, i.e. 5 to 10 centimeters an hour, makes it a highly specific symptom of strongyloidiasis [11, 13]. Finally, and although this aspect of infection needs to be confirmed, *S. stercoralis* infection might be associated with growth retardation in children [17]. Due to this combination of significant morbidity and high prevalence, *S. stercoralis* has been recognized as a public health problem in Cambodia. However, the national prevalence and the location of high risk zones are unknown.

A highly sensitive diagnostic approach consists in combining the Baermann and Koga agar plate culture techniques but this method is costly, time and labor consuming and requires laboratory staff specifically trained to identify *S. stercoralis* larvae by microscopy. Serological diagnosis is more sensitive than most coprological approaches but its use may be limited in endemic settings due to cross-reaction with other helminths species [18, 19]. Another issue is that serology may overestimate prevalence in endemic areas as it detects parasite-specific antibodies or antigens that can still be present long after contact with the parasite or cure, and cannot distinguish current from past infections [18]. While this last aspect would be an issue for cure assessment, it would not affect prevalence estimates in a population naïve to treatment against the investigated parasite. A serological test using an antigen from *S. ratti* to detect antibodies in urine was recently developed in Thailand [20, 21]. This technique has several strengths. While collecting urine samples is much easier than fecal samples, this test has a high sensitivity for *S. stercoralis* detection and does not cross-react with other soil-transmitted helminth (STH) species [20, 21].

Geostatistical models have been increasingly used in the past decade to delineate risk zones for helminthic infections at small and large scale and help targeting control efforts in areas of highest need [22–29]. Based on the association between environmental variables and infection levels at survey locations, such models can be used to predict infection levels through entire geographical zones.

The aim of this work was to estimate *S. stercoralis* prevalence in Cambodia and to predict *S. stercoralis* infection risk throughout the country to help guiding control efforts. A national parasitological survey was conducted in 2016 in all provinces of Cambodia to assess the infection with *S. stercoralis* based on a serological diagnosis using antigens of *S. ratti* [20]. Subsequently, geostatistical modeling was used to predict infection risk throughout the country.

## Methods

### Ethics statement

The study was approved by the National Ethics Committee for Health Research, Ministry of Health, Cambodia (NECHR, reference number 188, dated 02.05.2016). Prior to enrolment, all participants were explained the study goals and procedures. All participants aged 16 years and above provided written informed consent and parents or legal guardians provided consent for participants aged 6–15 years. All *S. stercoralis* cases were treated with a single oral dose of ivermectin (200µg/kg BW) and all other diagnosed parasitic infections were treated according to the national guidelines [30].

### Study setting

Cambodia counted 15.6 million inhabitants in 2015, 79.3% of whom lived in rural areas. [31]. The country has been undergoing fast economic development in the past decades, and with a Human Development Index rank of 143/188 in 2016 Cambodia belonged to the group of lower middle-income country in the World Bank classification [31, 32]. Although poverty strongly decreased in the past years and the proportion if the population living in extreme poverty was down to 2.2% in 2016, about one person in 5 (21.6%) still lived with less than 3.1 US$/day in 2016 [31]. Adult literacy and primary school net enrolment were 74% and 95%, respectively, in the years 2010-2014 and 32% of children aged below 59 months were stunted in 2015 [32]. Regarding water and sanitation in 2015, 42% and 69% of the rural population had access to improved sanitation facilities and improved water, respectively, while those figures were 88% and 100% for the urban population, respectively [32].

### Study population and design

A cross-sectional community-based survey was conducted among the general population in all 25 provinces of Cambodia between May and August 2016. In each province, 10 villages were randomly selected. Overall, eighteen villages originally selected were replaced because there remoteness compromised the quality of collected samples for parasitological data. In each village, households were selected using a systemic proportional sampling and all household members present on the survey day were enrolled until a maximum of 35 participants per village was reached. All household members aged 6 years and above were eligible. All *S. stercoralis* cases were treated with a single oral dose of ivermectin (200µg/kg BW) and all other diagnosed parasitic infections were treated according to the national guidelines [30].

### Assessment of *Strongyloides stercoralis* infection

Participants were asked to provide a urine sample on which *S. stercoralis* was diagnosed using an enzyme-linked immunosorbent assay (ELISA) based on *S. ratti* antigens [20]. After collection, urine specimens were preserved in NaN3 with the final concentration of 0.1% and kept at 4 °C at all until required for analysis. Samples were sent to the central laboratory of the National Centre for Parasitology, Entomology and Malaria Control (CNM) in Phnom Penh and from then sent to Khon Kaen University, Thailand, to proceed to the ELISA test. This method has shown to have no cross-reactivity with other STH, and has a high sensitivity, of 92.7%. *S. ratti* antigens may cross-react with filarial parasites, which are merely absent now from Cambodia, as well as with the liver fluke *Opisthorchis viverrini*, although very weakly [21, 33].

### Individual risk factor data

An individual questionnaire including demographics (age, sex, education attainment, main occupation), the number of household members, as well as access to sanitation (latrine availability at home, usual defecation place) and knowledge on worm infections (transmission route of and health problems caused by helminths) was administered to all study participants.

### Environmental data

Environmental parameters were extracted from freely available remote sensing (RS) sources for the period September 2015-August 2016, which corresponds to 1 year back from the last month of the study. Day and night land surface temperature (LST), international geosphere biosphere programme (IGBP) type 1 land use/land cover (LULC) as well as normalized difference vegetation index (NDVI) and enhanced vegetation index (EVI) were extracted at 1 x 1 km resolution from Moderate Resolution Imaging Spectroradiometer (MODIS) Land Processes Distributed Active Archive Center (LP DAAC), U.S. Geological Survey (USGS) Earth Resources Observation and Science (EROS) Center (http://lpdaac.usgs.gov). Rainfall data was obtained from WorldClim (www.worldclim.org). Digital elevation data were retrieved from the NASA Shuttle Radar Topographic Mission (SRTM) and CGIAR-CSI database, whereas distance to large water bodies was obtained from Health Mapper.

### Data management

Laboratory and questionnaire data were double-entered and validated in EpiData version 3.1 (EpiData Association; Odense, Denmark). Environmental data processing, geo-referencing and maps were done in ArcGIS version 10.2.1 (ESRI; Redlands, CA, United States). LULC 18 classes were merged into four categories according to similarity and respective frequencies. Year and seasonal means, maxima and minima of monthly means of EVI, LST and RFE were calculated and standardized. Environmental data were linked to parasitological and questionnaire data according to geo-referenced location. Data management and non-Bayesian data analysis were done in STATA version 13.0 (StataCorp LP; College Station, United States of America). Bayesian geostatistical models were fitted using WinBUGS version 1.4.3 (Imperial College & Medical Research Council; London, UK). Age was grouped in five classes, as follows: (i) 6-12 years, (ii) 13-18 years, (iii) 19-30 years, (iv) 31-50 years, and (v) >50 years. Predictions at un-surveyed locations were performed in Fortran 95 (Compaq Visual Fortran Professional version 6.6.0, Compaq Computer Corporation; Houston, United States of America).

### Statistical Analysis

Chi-square (χ^2^) test was used to compare proportions. The association between infection risk and covariates was assessed, using mixed non spatial bivariate logistic regressions accounting for village clustering, i.e. with a non-spatial village-level random effect. Covariates exhibiting an association at a significance level of at least 15%, as determined by the likelihood ratio test (LRT), were included in the multivariate logistic regression models. In case of correlated variables, the variable resulting in the model with the smallest Akaike’s information criterion (AIC) was selected. For the risk factor analysis, variables exhibiting high Wald p-values were removed one by one and kept outside the model if their removal resulted in a lower AIC. Summary measures of continuous environmental variables, i.e. LST day and night, rainfall and distance to water were standardized before inclusion in the multiple regression models. To explore the relationship between *S. stercoralis* infection risk and age, smoothed age-prevalence curves were produced with the “mkspline” command in STATA that regresses each outcome against a new age variable containing a restricted cubic spline of age.

For geostatistical models, a stationary isotropic process was assumed, with village-specific random effects following a normal distribution with mean zero and a variance-covariance matrix being an exponential function of the distance between pairs of locations. Vague prior distributions were chosen for all parameters. Further information on model specification is available in S1 Appendix. Markov chain Monte Carlo (MCMC) simulation was used to estimate model parameters [34]. Geostatistical models were run using the WinBUGS “spatial.unipred” function [35]. Convergence was assessed by examining the ergodic averages of selected parameters. For all models, a burn-in of 5,000 was followed by 30,000 iterations, after which convergence was reached. Results were withdrawn for the last 10,000 iterations of each chain, with a thinning of 10. Model fit was appraised with the Deviance Information Criterion (DIC). A lower DIC indicates a better model [36].

Three types of Bayesian mixed logistic models were run. First, models without covariates using alternately a geostatistical or an exchangeable random effect were run to quantify the extent of village-level spatial correlation and unexplained variance of *S. stercoralis* prevalence. Second, a risk factor analysis model was used to assess individual-level demographic, sanitation, and knowledge risk factors, as well as environmental covariates associated with infection risk. Third, a model including only environmental covariates was aimed at predicting infection risk at non-surveyed locations.

### Prediction of *S. stercoralis* at non-surveyed locations

For model validation, 199 (80%) randomly selected villages were used for fitting and the 50 (20%) remaining were used as test locations. A pair of models containing the same covariates but including alternately a non-spatial (exchangeable) or spatial (geostatistical) random effect was run. Model predictive ability was assessed by comparing the Mean Squared Error (MSE), which is obtained by squaring the average of absolute differences between predicted and observed prevalence rates at test locations.

Using the model with the best predictive ability, *S. stercoralis* infection risk was predicted at 68,410 pixels of 2×2 km resolution using Bayesian Kriging [37].

## Results

### Study population

Among the 8,661 participants enrolled in the study, 1407 did not provide any urine and 338 were discarded because they did not provide stool sample –which was requested for other assessments not presented in this work-, 8 participants did not have any questionnaire data. Overall 7,246 participants living in 2,585 households and 249 villages were included in the analysis. The mean number of participants per village was 30.2 with an interquartile range of 6, and a minimum of 5. Except the villages Ou Tracheak Chet in Preah Sihanouk Province (5 participants) and Kampong Chrey in Preah Vihear province (9 participants), all villages had more than 10 participants and 93.6% of villages had 20 participants or more. Table 1 shows the characteristics of participants with complete parasitological and questionnaire data.

**Table 1.**
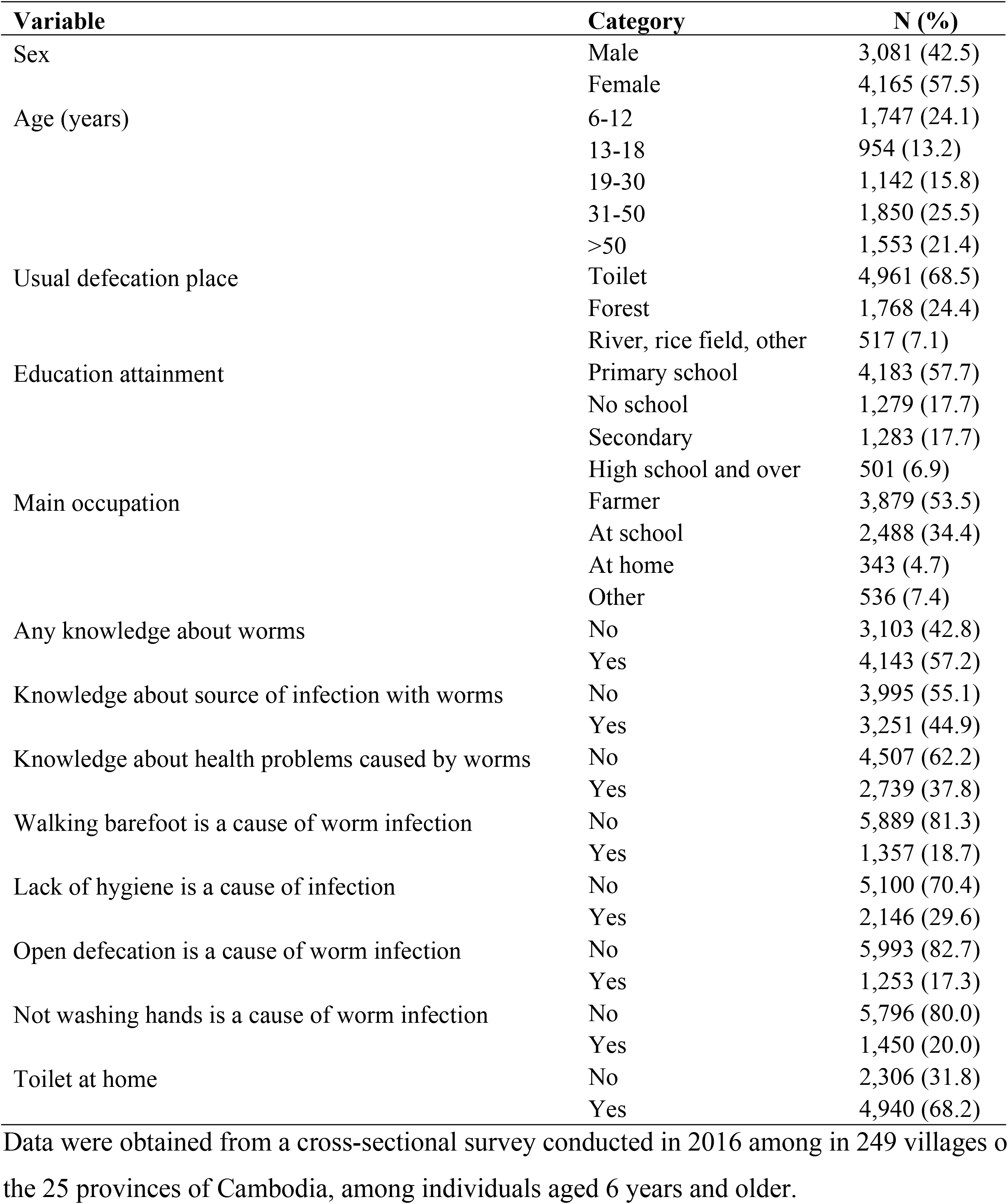
Characteristics of the 7,246 participants included in the analysis

Females (57.5%) were overrepresented in the sample compared to their proportion in the Cambodian population (51.5%) as assessed by the 2013 inter-census population survey [38]. The age distribution of the sample was very similar to that of the total Cambodian population: children and adolescents aged up to 14 years represented 29.95% and 29.4%, adolescents and adults aged 15 to 64 years adults represented 65.6% and 64.2%, and elderly adults aged 65 and above represented 5.8% and 5.0% of the sample and the Cambodian population, respectively.

The proportion of males and females were similar in participants excluded or included in the analyzed sample, while children and young adults aged between 6 and 30 years were less represented (53.0%) in the sample than among excluded participants 64.3%). Similarly, farmers were overrepresented (53.6% of the sample *vs*. 41.1% of excluded participants) and scholars underrepresented (34.3% of the sample *vs*. 51.6% of excluded participants) in the final sample. There was no difference between participants excluded from, or included in, the analyzed sample in terms of usual defecation place.

### *Strongyloides stercoralis* prevalence

Overall, *S. stercoralis* prevalence was 30.7% (95% confidence interval (CI): 29.7 – 31.8), ranging at province level from 10.9% (95%CI: 7.4 – 14-4) in Prey Veng province to 48.2% (95%CI: 42.2 – 54.1) in Koh Kong province. Fig 1 shows the provinces of Cambodia and Fig 2 displays province level prevalence rates. Prevalence was highly variable at village level. The smallest prevalence rate, of 2.9% (95% CI: 0.1 – 14.9), was found in a village of Kandal province, where only 1 of 35 participants was infected. The highest rates were 88.9% (95%CI: 51.8 – 99.7) and 80% (95%CI: 63.1 – 91.6), observed in one village in Preah Vihear province and another village in Koh Kong province, respectively. However there were only 9 participants in the Preah Vihear province village across provinces. The map presented in Fig.3 displays observed *S. stercoralis* prevalence in each surveyed village.

**Figure 1.**
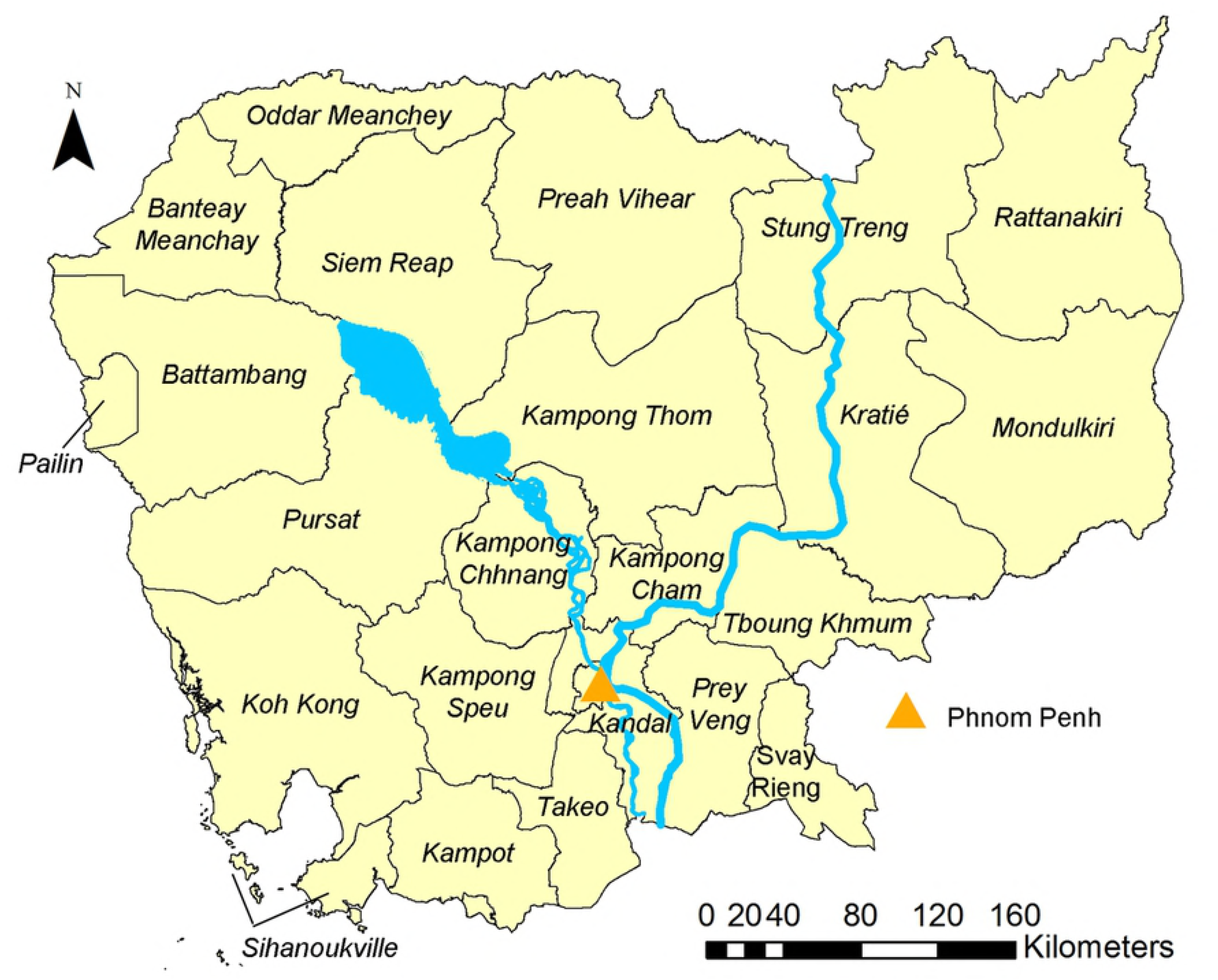
Map of Cambodian provinces. This map was created with ArcGIS version 10.0 (ESRI; Redlands, CA, USA) specifically for this study by Forrer et al.

**Figure 2.**
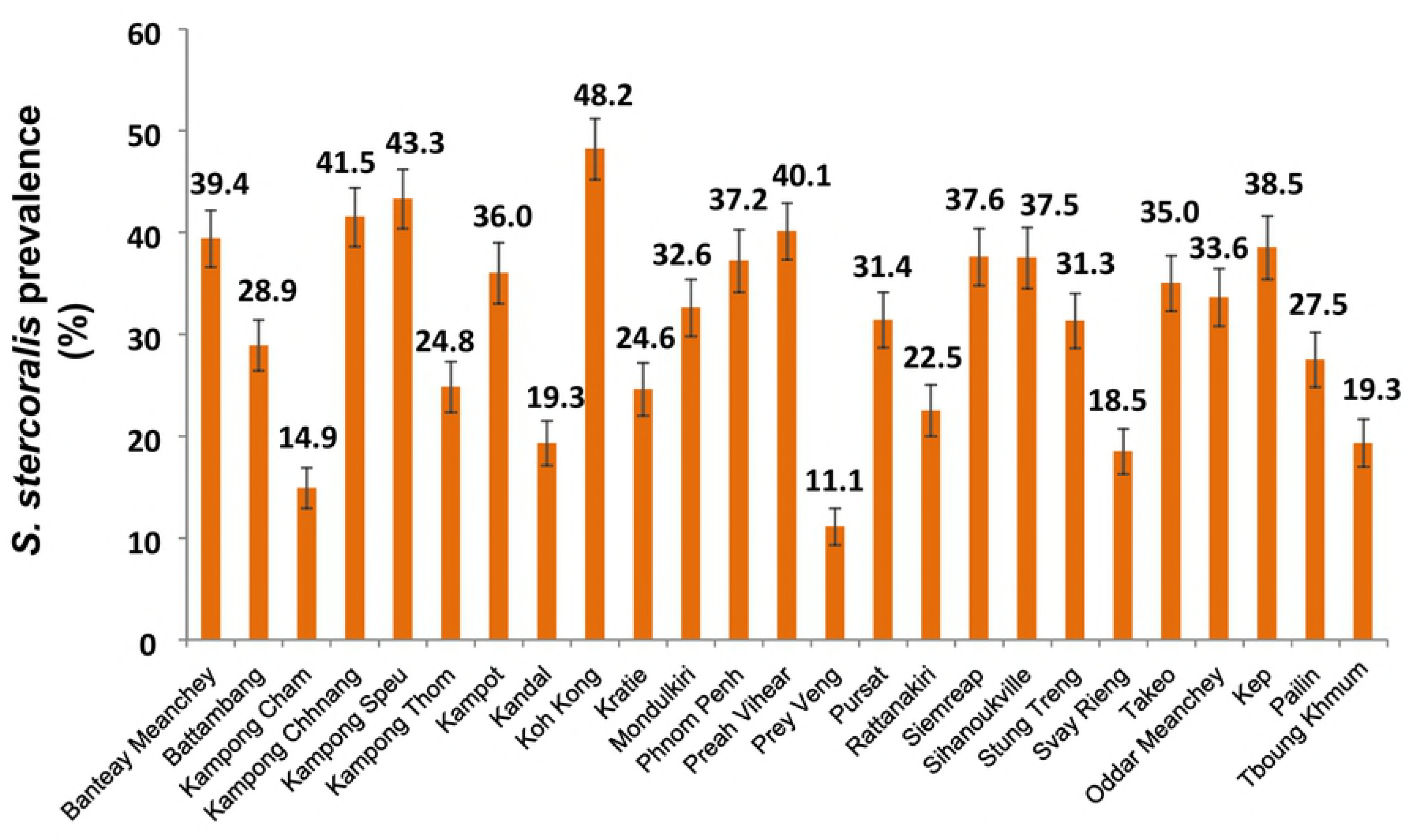
Province-level *S. stercoralis* prevalence in 25 provinces of Cambodia Data were obtained from a cross-sectional survey conducted in 2016 in 249 villages of Cambodia, among 7,246 participants aged 6 years and above.

**Figure 3.**
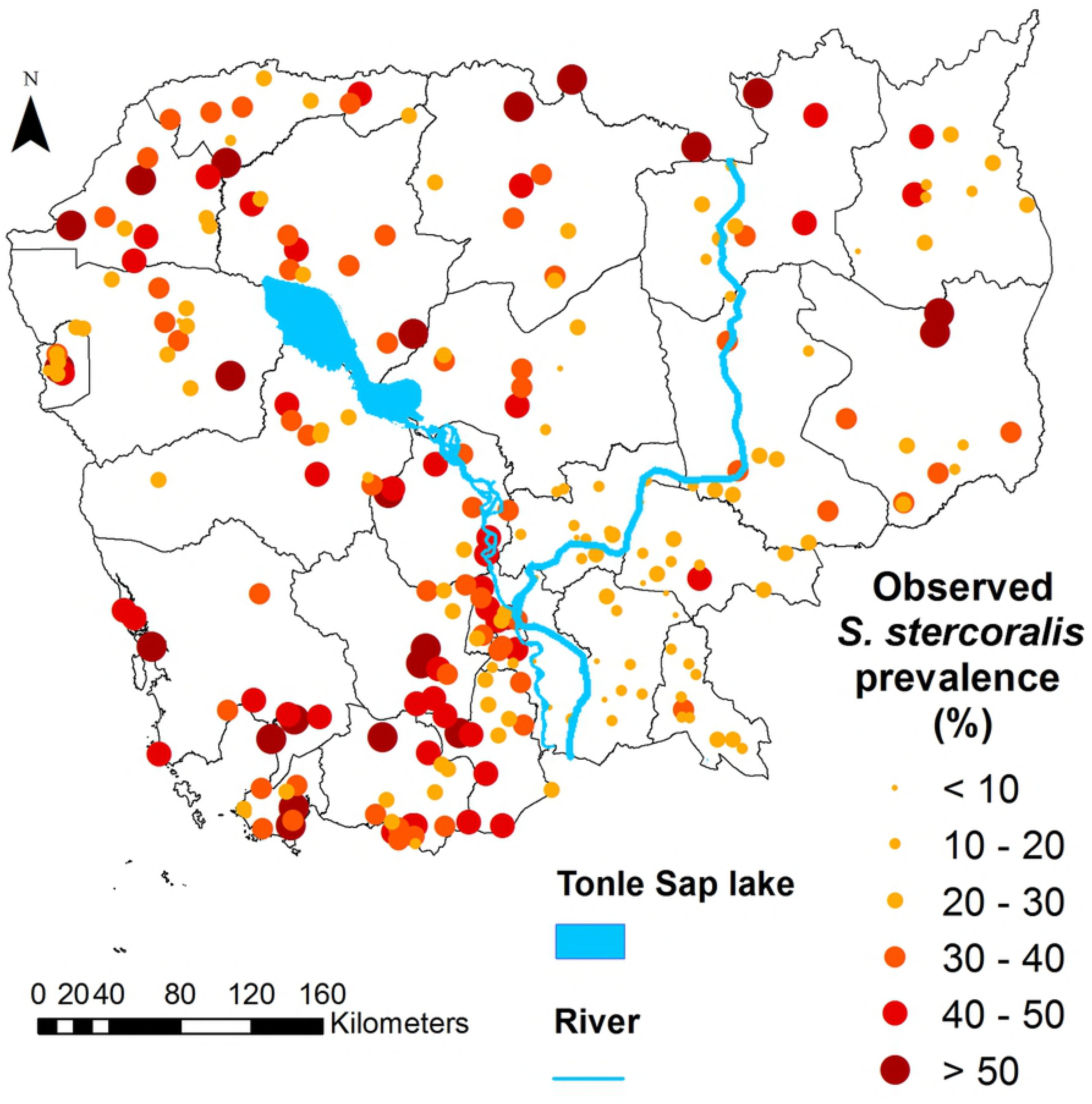
Map of Cambodia showing observed *S. stercoralis* prevalence in the 249 study villages, Cambodia. Data were obtained from a cross-sectional survey conducted in 2016 in 249 villages of Cambodia, among 7,246 participants aged 6 years and above. This map was created with ArcGIS version 10.0 (ESRI; Redlands, CA, USA) and display the results obtained specifically from this study by Forrer et al.

### Spatial correlation

The model parameters of three geostatistical models, i.e. (i) model without covariates, (ii) the predictive model including only environmental variables, and (iii) the risk factor analysis model including environmental, demographic and behavioral covariates are presented in Table 2. In absence of explanatory variables, *S. stercoralis* risk clustered at a distance of 85 km (range). Most of *S. stercoralis* tendency was due to environmental covariates as indicated by the range of 3.2 km after introducing environmental variables (predictive model).

**Table 2.** Parameters of three geostatistical models

|  | No covariates <sup>(a)</sup> |  | Predictive model <sup>(b)</sup> |  | Risk factor analysis model <sup>(c)</sup> |  |
| --- | --- | --- | --- | --- | --- | --- |
|  | median | 95% BCI | median | 95% BCI | median | 95% BCI |
| Range (km) | 85.3 | 1.10 - 185.8 | 3.20 | 1.10 - 99.4 | 2.80 | 1.10 - 49.7 |
| $\rho$ | 3.8 | 1.78 - 240.0 | 105.60 | 3.35 - 282.6 | 116.10 | 6.55 - 283.60 |
| $\sigma^2$ | 0.36 | 0.21 - 0.59 | 0.27 | 0.19 - 0.40 | 0.29 | 0.21 - 0.41 |
| DIC |  |  |  |  | 8180.98 | - |

$\sigma^2$ is the location-specific unexplained variance.
$\rho$ is the decay parameter. The range (range=3/ $\rho$ ) is the distance at which the spatial correlation becomes less than 5%.
<sup>(a)</sup> Geostatistical model without covariates
<sup>(b)</sup> Predictive model: with environmental covariates only
<sup>(c)</sup> Risk factor analysis model: with environmental and demographic and behavioral covariates

### Result of the model validation and predictive model

The predictive ability of the geostatistical model (MSE=182.9, DIC=6894.3) including environmental covariates (predictive model) was slightly higher than that of its non-spatial counterpart (MSE = 187.7, DIC = 6894.4). Therefore the geostatistical model was used to predict *S. stercoralis* risk at non-surveyed locations. The geographical distributions of the covariates used in the geostatistical predictive model, together with elevation, are displayed in S1 Figure. Odds ratios of those covariates are presented in Table 3.

**Table 3.** Results of the geostatistical predictive model

| Variable | Category | OR | 95% BCI |
| --- | --- | --- | --- |
| LST night dry season maximum (°C) | - | 1.21 | 1.05 - 1.33 |
| Rainfall year minimum (mm/month) | - | 1.35 | 1.10 - 1.49 |
| Distance to water (km) | - | 1.10 | 0.97 - 1.22 |
| Land use, land cover | Crops & natural vegetation mosaic, grass | 1 | - |
|  | Cropland | 0.82 | 0.67 - 1.03 |
|  | Forest and savanna | 1.17 | 0.89 - 1.52 |
|  | Water and wetlands | 1.29 | 0.88 - 1.91 |
| Range (km) | - | 3.20 | 1.10 - 99.4 |
| $\rho$ | - | 105.6 | 3.35 - 282.6 |
| $\sigma^2$ | - | 0.27 | 0.19 - 0.40 |

LST: Land surface temperature; BCI, Bayesian credible interval; OR: odds ratio; OR in bold are significant at 5% level $\sigma^2$ is the location-specific unexplained variance. $\rho$ is the decay parameter. The range (range=3/ $\rho$ ) is the distance at which the spatial correlation becomes less than 5%.

### Risk factors for *S. stercoralis* infection

The results of non-spatial bivariate mixed regressions are presented in S2 Table. The results of the multivariate Bayesian geostatistical risk factor analysis are presented in Table 4.

**Table 4.**
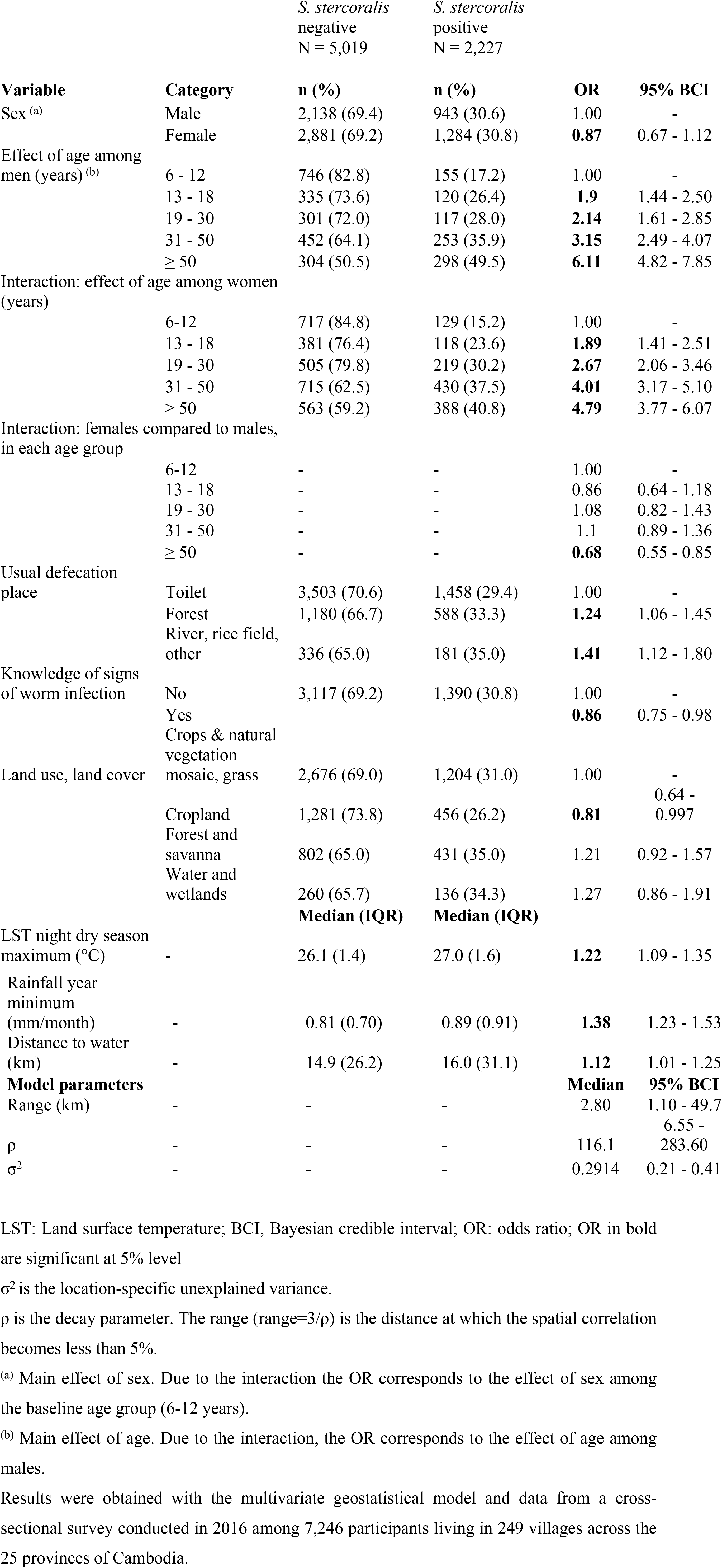
Results of the risk factor analysis

Sex was an effect modifier of age. Infection risk increased with age for both genders but women aged 50 years and above had a lower risk of being infected than males. The relationship between *S. stercoralis* infection risk and age is presented in Figure 4. Participants usually practicing open defecation (31.5% of participants defecating either in forests, or rice field and water) had higher odds of being infected, while individuals who had some knowledge about health problems resulting from worm infection has lower odds of harboring *S. stercoralis*. As for environmental factors, *S. stercoralis* infection risk was positively associated with increasing night land surface temperature (LST night) dry season maximum, increasing minimum year rainfall and increasing distance to water. Finally, the odds of *S. stercoralis* infection were lower among participants living in villages located in croplands, i.e. rice fields.

**Figure 4.**
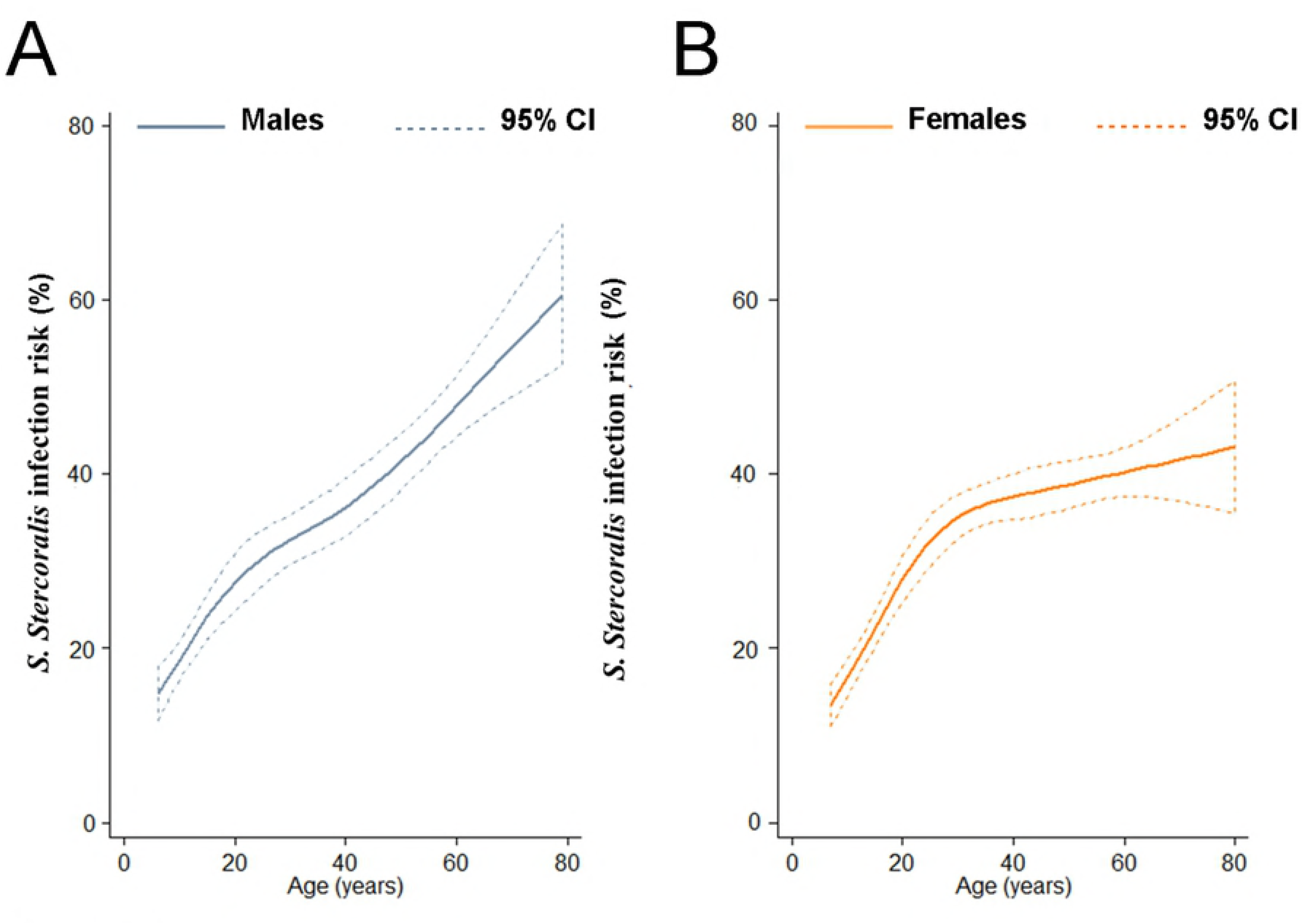
Smoothed age-prevalence of *S. stercoralis*, Cambodia Data were obtained from a cross-sectional survey conducted in 2016 in 249 villages of Cambodia, among 7,246 participants aged 6 years and above. Restricted cubic splines were used. Data are stratified for males (A) and females (B). Uncertainty is expressed as 95% confidence interval (CI).

### Spatial prediction of *S. stercoralis* infection risk

Figure 5 and 6 display the predicted median *S. stercoralis* prevalence in Cambodia and the lower and upper estimates, respectively. Prevalence was consistently higher than 10% except in a small area of Prey Veng province. *S. stercoralis* predicted risk was below 20% only in five provinces, i.e. Kampong Cham, Tboung Khmum, Prey Veng, Kandal and Svay Rieng. Predicted prevalence was particularly high in the north of Preah Vihear and Stung Treng provinces near the Lao border, as well as in the South, in areas of Kampong Speu, Koh Kong, Preah Sihanouk, and Kampot provinces.

## Discussion

We present here, to our knowledge, the first national prevalence estimate and nationwide infection risk map of *S. stercoralis*. The infection is ubiquitous in Cambodia. Based on a sample encompassing all the 25 country provinces and including over 7,200 participants, *S. stercoralis* occurs in Cambodia at prevalence rates systematically over 10%, with a national prevalence of 30%.

Infection risk was the lowest in the Southeast of the country, i.e. in the provinces of Prey Veng, Kandal, Kampong Cham, as well as the West and South parts of Tboung Khmum and Kampong Thom provinces, respectively. The highest province-level prevalence rates, above 40%, were found in Preah Vihear in the North, Kampong Chhnang in the Centre and in the South, Koh Kong and Kampong Speu.

The size of *S. stercoralis* infection clusters was relatively small, 85 km, similarly to that observed for hookworm infection risk in the country [25]. Almost all spatial correlation of *S. stercoralis* infection was explained by its association with environmental factors (as indicated by the dramatic drop of the range, down to 3.2 km, after introducing environmental covariates in the model). This result is not surprising as in absence of available treatment the parasite biological requirements would mostly condition its distribution. The distribution of hookworm prevalence among school-aged children in Cambodia was similar to that of *S. stercoralis*, likely due to the resembling transmission routes of those two nematodes. Yet, the area with lower hookworm prevalence was larger, probably because of ongoing STH deworming programs [25].

The odds of being infected increased with increasing maximum night temperature and increasing minimum rainfall. In presence of sufficient humidity hookworm has a good tolerance for high temperatures with its larvae having the ability to migrate in the soil, and *S. stercoralis* larvae might have the same ability [39]. The positive association between temperature and risk is more surprising, although this might relate to a particularity of *S. stercoralis* life cycle. The number of females and infective larvae developing in the external environment depends on temperature, with numbers of infective larvae being maximum when temperatures are of 30°C and above [11]. Hence, night maximum temperatures which range between 24°C and 32°C in Cambodia might have an impact on the amount of infective larvae present in the environment.

Regarding the environmental predictors of *S. stercoralis* infection, distance to water and the land cover category of cropland were not significantly associated with infection risk in the predictive model but became significant in the risk factor analysis after adjusting for demographic and behavioral factors. We found a positive association between *S. stercoralis* infection risk and distance to water. The development and survival of *S. stercoralis* larvae is affected by immersion, so seasonal flooding might affect their survival in areas close to water bodies [40, 41]. Similarly, this relationship between larvae survival and water might explain the lower infection in areas occupied by croplands, which mostly correspond to rice fields that are regularly flooded. Yet it is also possible that distance to water captured other unmeasured features, including factors relating to socio-economic features and human activity [27]. In Cambodia, people have a clear preference for pour-flush latrines and would rather not have a toilet than pit latrines, but pour flush latrines function only with water [42]. Limited availability of water due to living away from permanent water bodies might result in decreased access to, or use of, sanitation facilities.

Studies that investigated risk factors for *S. stercoralis* infection mostly report a higher risk in men, whereas the relationship between age and *S. stercoralis* prevalence seems to vary across settings [9, 10, 43, 44]. In this national survey among more than 7,200 individuals aged six years and above, we found that prevalence increased with age both in men and women, although men aged 50 years and above had higher odds of *S. stercoralis* infection than women of the same age. Boys aged between 6 and 12 years were also more likely to harbor *S. stercoralis* than their female counterparts. Previously to this national survey, prevalence was found to increase with age and reach a plateau in adulthood in North Cambodia, while no cases were found in individuals aged less than 15 years in Yunnan, China [9, 45, 46]. Yet, no association between age and *S. stercoralis* infection was found in Lao PDR, South Cambodia, or Zanzibar [10, 44, 47]. Age-specific infection risk is of particular importance to target control programs and should be further documented.

An interesting finding was that individuals who declared having some knowledge about health problems caused by worm infections had lower odds of being infected with *S. stercoralis*, but having some knowledge about sources of infection was not associated with infection risk. While knowledge does not necessarily translate into behavior change, this result suggests that being aware of personal disease risk, –which is an important driver in health promotion and increases compliance to helminth control programs– might be a better trigger of hygienic behavior than knowing exposure sources [48, 49].

The protective effect of improved sanitation against STH infection is widely acknowledged [50–54]. We found that, compared to open defecation, defecating in latrines was protective against *S. stercoralis* infection. This result is in line with other studies conducted in Cambodia or Ecuador, and with a recent meta-analysis that included 9 studies investigating the impact of sanitation on *S. stercoralis* infection risk, and estimated a pooled OR of 0.50 (95%CI: 0.36-0.70) [9, 10, 54–57]. Of note, village-level sanitation coverage was also found to reduce re-infection risk one year after treatment in North Cambodia [46].

The present work has several limitations. First, women were overrepresented in the sample compared to the general Cambodian population and the lower prevalence among young girls and women aged 50 years and above, compared to males, might have resulted in a slight underestimation of prevalence. Most importantly, our sample was representative of the 2013 Cambodian general population in terms of age, which is strongly associated with infection risk [38].

Second, it was the first time that the serological diagnostic method of detecting IgG anti-bodies employed in this study was used for a large-scale survey. This method has proven a high sensitivity for *S. stercoralis* detection, and it does not suffer from cross-reactivity with other STH [20, 21]. However, validation of the method in different settings should be carried out. Finally, our risk factor analysis did not adjust for socio-economic status which was found associated with infection risk in North Cambodia, but results from the few studies that accounted for it are heterogeneous [9, 10, 46, 58, 59]. Given the strong association between poverty and other STH infections, it is likely that *S. stercoralis* risk distribution is also associated with socioeconomic status and future studies should account for it. Of note, the socioeconomic status was not a confounder of the relationship between age or sex and *S. stercoralis* infection risk in North Cambodia and would probably not have substantially affected the estimates for sex and age in the present study [46, 59].

Our study represents a clear risk map of *S. stercoralis* of a highly endemic setting. Based on these data the population at risk can be quantified and planning of concrete control approach become realistic. Further developing this operational approach in other settings and with further validated diagnostic approaches will result in data bases for global planning. The mainstay of the WHO strategy to control soil-transmitted helminths is preventive chemotherapy, i.e. the regular treatment with mebendazole or albendazole to prevent high intensity infections and the associated morbidity to entire populations or at-risk groups [60, 61]. However those drugs are not efficacious at a single oral dose against *S. stercoralis* for which the drug of choice is ivermectin [62–64]. A single oral dose (200µg/kg Body Weight) of ivermectin has recently been found to achieve a high cure rate and result in re-infection rates below 15%, one year after treatment in a highly endemic setting of Cambodia [46, 62, 63]. As our results demonstrate, *S. stercoralis* is highly endemic in all provinces in Cambodia and the inclusion of ivermectin in the control program would be required [13, 64, 65]. Yet, this drug is not subsidized in in regions where onchocerciasis is absent, let alone to treat *S. stercoralis*, and its high cost in Cambodia, 10 USD per tablet –while up to 5 tablets may be needed to treat an individual, depending on their weight–, precludes the deployment of control measures in the country.

In absence of data on age specific morbidity, the fact that individuals of any age appear to have the same risk for reinfection one year after treatment suggests community-wide control [46]. Yet, it appeared in a study investigating *S. stercoralis* related morbidity in Cambodia that children and adolescents with higher parasite loads had higher odds of being stunted, while *S. stercoralis* infection was found to be associated with anemia but not stunting in Argentina [17, 66]. The relationship between *S. stercoralis* parasite loads, morbidity and transmission intensity needs to be assessed, as well as age-related infection levels, using appropriately designed longitudinal studies. Cost-effectiveness studies of various control options are needed while mathematical models could help better appraising the parasite transmission dynamics and guide control efforts, as the complex life cycle of *S. stercoralis* life might yield transmission dynamics that are different from the other STH.

Cambodia benefits from a well-established STH control network and was among the first countries to reach the 75% national coverage target [67, 68]. STH control was recently scaled up to reach children in secondary and high schools, including in private schools, and women of child bearing age in factories [69]. Additionally, schistosomiasis has been successfully controlled with no severe cases recorded recently and lymphatic filariasis has been eliminated as a public health problem and is now under surveillance for elimination [68, 70, 71].

However, in this country that has demonstrated its capacity to efficiently address helminthic infections, the control of *S. stercoralis* is currently hindered by the high cost of ivermectin, which cannot be entirely supported by the Ministry of Health. Either subsidizations, donations, or the production of affordable generics are necessary to start tackling this dangerous parasite that infects almost a third of the Cambodian population.

## Acknowledgements

We are grateful to all of the study participants. Our sincere thanks go to the laboratory technicians and staff at the Helminth Control Program of the National Centre for Parasitology, Entomology and Malaria Control, Phnom Penh, Cambodia, and the staff of the laboratory staff of the Parasitology department of the Khon Kaen University, Khon Kaen, Thailand. We thank the Provincial Health Departments of all provinces for their support and field work, and the local authorities for their support.

## Supporting information captions

**Table S1:** Results of the bivariate non-spatial regression for individual-level risk factors

**Figure S1:** Environmental predictors

**Appendix S1:** Baysian model formulation

**STROBE** Strobe statement – checklist of items that should be included in reports of cross-sectional studies

